# Defective linear and circular RNAs biogenesis in Huntington’s disease: CAG repeat expansion hijacks neuronal splicing

**DOI:** 10.1101/2021.12.27.474266

**Authors:** Dilara Ayyildiz, Alan Monziani, Takshashila Tripathi, Jessica Döring, Guendalina Bergonzoni, Emanuela Kerschbamer, Francesca Di Leva, Elia Pennati, Luisa Donini, Marina Kovalenko, Jacopo Zasso, Luciano Conti, Vanessa C. Wheeler, Christoph Dieterich, Silvano Piazza, Erik Dassi, Marta Biagioli

## Abstract

Alternative splicing (AS) appears to be altered in Huntington’s disease (HD), but its significance for early, pre-symptomatic disease stages has not been inspected.

Here, taking advantage of *Htt* CAG knock-in mouse *in vitro* and *in vivo* models, we demonstrate a strong correlation between *Htt* CAG repeat length and increased aberrant linear AS, specifically affecting neural progenitors and, *in vivo,* the striatum prior to overt behavioral phenotypes stages. Remarkably, expanded *Htt* CAG repeats reflect on a previously neglected, global impairment of back-splicing, leading to decreased circular RNAs production in neural progenitors.

Though the mechanisms of this dysregulation remain uncertain, our study unveils network of transcriptionally altered micro-RNAs and RNA-binding proteins (CELF, hnRNPS, PTBP, SRSF) which, in turn, might influence the AS machinery, primarily in neural cells.

We suggest that this unbalanced expression of linear and circular RNAs might result in altered neural fitness, contributing to HD striatal vulnerability.

## INTRODUCTION

Huntington’s disease (HD) is a hereditary, fatal neurodegenerative disorder caused by a CAG trinucleotide expansion within exon 1 of the *HTT* gene (1993). Explicit clinical onset typically occurs in mid-life and leads to an inexorable decline to death after 10-15 years (Perutz et al. 1999). A polymorphic CAG tract up to 35 repeats is found in unaffected individuals, whereas alleles bearing 36 or more repeats lead to HD symptoms. Since the *HTT* gene is ubiquitously expressed during human development and in all body districts, the effects of the mutation is strongly pleiotropic (Bassi et al. 2017). The central nervous system, however, remains the main region affected by mutant *HTT*, with a prominent loss of GABAergic medium-sized spiny neurons of the striatum, constituting the major contributor to movement, cognitive and behavioral dysfunctions (Vonsattel et al. 1985; Rosas et al. 2003; Han et al. 2010). So far, the rate-limiting mechanism (s) of neurodegeneration remains elusive although chromatin, transcription, and RNA processing dysregulations are emerging as fundamental features (Bassi et al. 2017; Kerschbamer and Biagioli 2016). In particular, RNA processing and alternative splicing (AS) alterations might affect the level and composition of a broad repertoire of proteins, thus contributing to HD striatal vulnerability and pathogenesis. The spliceosomal activity can be directly modulated by the expression level and/or sequestration of various proteins that bind to nascent mRNA. Interestingly, huntingtin can associate with the WW-containing proteins HYPA and HYPC (*Htt* Yeast two-hybrid Protein A and C), also known as FBP11/PRPF40A, and PRPF40B, respectively (Faber et al. 1998; Jiang et al. 2011; Passani et al. 2000), participating in early spliceosomal assembly and 5’ site recognition. On the other hand, mis-splicing events in individuals with highly expanded *HTT* CAG repeats have been shown to produce the small, highly pathogenic, polyadenylated exon 1-intron 1 *HTT* transcript (Neueder et al. 2017). Moreover, tissue-restricted trans-splicing regulators, binding auxiliary exonic and intronic cis-regulatory signals, modulate splice-site choice by interacting with components of the splicing machinery (Wahl et al. 2009; Ule and Blencowe 2019). Recent evidence supports a role for huntingtin in regulating the expression of four RNA-binding proteins (PTBP1, SFRS4, RBM4, SREK1) in HD *post-mortem* brains, thus, in turn, affecting the relative abundance of specific target isoforms (Lin et al. 2016). A third layer of complexity in AS regulation is added by broad chromatin conformation and transcriptional kinetics (Andersson et al. 2009; Spies et al. 2009; Schwartz et al. 2009). In fact, chromatin relaxation accelerates RNA Pol II processing, correlating with alternative exon skipping. Conversely, packed nucleosomes slow down RNA Pol II progression, causing a pause in transcription and not-constitutive weak exon inclusion. Interestingly, altered chromatin remodeling and histone modifications enrichment emerge as key features in HD. The work from Vashishtha et al., 2013 shows changes in H3K4me3 in HD mice and *post-mortem* tissues (Vashishtha et al. 2013), whereas we recently described an effect of wild-type and mutant huntingtin on PRC2 and Mll-containing complexes (Biagioli et al. 2015; Seong et al. 2010). In addition, huntingtin can directly interact with the HypB/SetD2 H3K36me3 methyltransferase, implicated in RNA splicing and Pol II elongation of transcription (Faber et al. 1998; Passani et al. 2000; Simon et al. 2014; Zhu et al. 2017). As experimental evidence points toward dysregulated AS as a key feature in HD pathology, it is tempting to hypothesize that nuclear huntingtin might regulate AS outcome through different direct and/or indirect modes, thus contributing to HD pathogenesis.

Importantly, AS regulation is crucial not only to the establishment of a repertoire of protein-coding isoforms extremely relevant for the proper physiology of the nervous system, but also to the biogenesis of circular RNAs (circRNAs), unusually stable non-coding RNAs produced by the circularization of exons (Jeck et al. 2013; Memczak et al. 2013; Salzman et al. 2012). CircRNAs are highly enriched in neurons and have been implicated in a wide variety of pathological conditions, including neurological diseases (Holdt et al. 2018). CircRNA biogenesis seems to correlate with exon-skipping events, but the exact mechanisms of spliced exon-exon circularization remains unclear (Kelly et al. 2015; Barrett et al. 2015). The vast majority of circular RNAs localize in the cytoplasm and their exonic sequences might encompass micro RNAs (miRNAs) and RNA Binding Proteins (RBPs) binding sites (Memczak et al. 2013; Hansen et al. 2013), thus acting as competing-endogenous RNA (Salmena et al. 2011). In this scenario, a circRNA could efficiently engage (“sponge”) either miRNAs or RBPs, or both, eventually relieving their canonical mRNA targets from post-transcriptional regulation (Memczak et al. 2013; Piwecka et al. 2017; Hansen et al. 2011; Abdelmohsen et al. 2017; Chen et al. 2019). Whether mutant huntingtin may modulate circular RNA expression levels remains unexplored.

Here, we investigated whether linear and back-splicing processes, producing linear mRNAs and circular isoforms, might be altered by the HD mutation in a CAG-dependent and tissue-specific manner. We unveiled new mechanisms dysregulated in HD at the genome-wide level and possible regulatory cross talks with miRNAs and RBPs.

## RESULTS

### Aberrant linear alternative splicing shows *Htt-*CAG length and age dependency, specifically in the mouse striatum

In order to gauge evidence of AS alterations in the brain regions ofneural HD animal models, we examined publicly available RNA sequencing (RNAseq) datasets (striatum (GSE65774), cortex (GSE65770) and liver (GSE65772), available through HDinHD, (see also Materials and Methods) (Langfelder et al. 2016). The AS events in striatum, cortex and liver of 6 knock-in (KI) mouse models (Q20, Q80, Q92, Q111, Q140 and Q175) carrying different *Htt* CAG repeat lengths were determined using rMATS (Shen et al. 2012, 2014; Park et al. 2013). Q20 mice, with the lowest *Htt* CAG repeat length, were used as controls. The total number of genes showing one of the five major types of AS events [alternative 5’ splice site (A5SS), alternative 3’ splice site (A3SS), mutually exclusive exons (MXE), retained intron (RI) and skipped exon (SE)] was evaluated using a collection of both reads spanning splicing junctions (JC) and reads spanning splicing junctions plus on target (JCEC) (Supplemental Table_S1).

The striatum, the most severely affected brain region in HD and HD model mice, exhibited the highest number of aberrantly spliced transcripts, while cortex and liver did not show overt aberrant splicing phenotypes (Fig. 1a, Supplemental_Fig_S1, Supplemental Table_S1). Strikingly, the alteration in the total number of mis-spliced events in the striatum strongly correlated (0.8 to 0.97 R^2^, Supplemental_Fig_S2) with *Htt* CAG length (Supplemental_Fig_S2), with Q175 mice showing the greatest number of abnormal splicing events at all three time points analyzed (2, 6 and 10 months). Cortical mis-splicing showed some, more variable, degree of CAG correlation at 2 and 6 months (∼ 0.6 to 0.9 R^2^, Supplemental_Fig_S2), while no significant association was detected in the liver. Interestingly, a *Htt* CAG correlated increase in aberrant striatal AS was already visible at early stages (2 months), becoming more significant as the pathologic process progressed (6-10 months) (Fig. 1a, Supplemental Table_S1).

**Figure 1.**
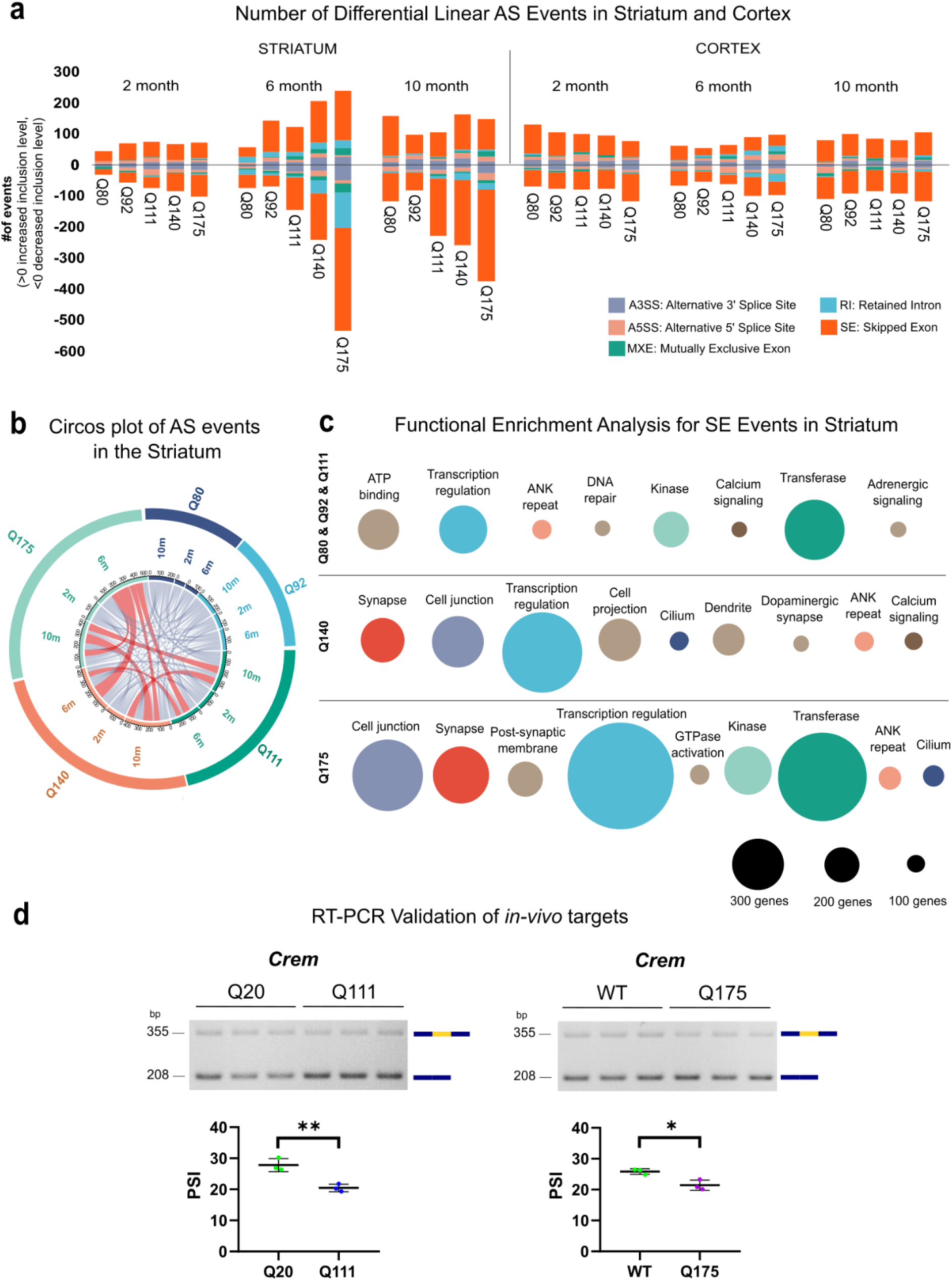
Aberrant linear alternative splicing in the striatum of KI animal models of HD. **a)** Bar graphs show the number of differential AS events in the striatum and cortex from 6 mouse KI models (Q80, Q92, Q111, Q140 and Q175) of HD, presenting different *Htt-* CAG repeat lengths and 3 ages (2, 6, and 10 months). The number of events is shown for each genotype, time point and brain region. The inclusion level is calculated in comparison to Q20 controls and the positive or negative values are plotted. Source data by Langfelder P. *et al* (2016). Further details can be found in the Methods section. Each color of the bar chart represents a different AS event. **b)** Circos plot represents the number of transcripts within the striatum - showing differential AS events - shared between different genotypes (Q80, Q92, Q111, Q140 and Q175) and time points (2, 6, and 10 months). Conditions (genotypes and/or time points) sharing more than 50 transcripts are depicted in red. **c)** Weighted nodes graphical representation shows the functional enrichment analysis for transcripts showing significant skipped exon (SE) events in the striatum. Highly expanded *Htt* CAG sizes (Q140 and Q175), the major contributors to aberrant SE in the striatum, are shown separated. Nodes’ size legend is depicted at the bottom. Nodes are ordered by adjusted p-values using Benjamini Hochberg (<0.05). The same enrichment terms shared between genotypes are presented with nodes of the same color, while unique terms are colored in light brown. **d)** Representative agarose gel images (top) and dot-plots (bottom) show the RT-PCR results of AS validation for *Crem*, a selected transcript target. RT-PCR assay and quantification were performed on RNA from striata from an independent set of wild-type (WT), Q20 and Q111/Q175 mice. *Crem* isoforms with inclusion or exclusion of the variable exon (in yellow) are visualized and quantified. Dot plots report the PSI, percent-spliced-in. *P < 0.05, **P < 0.01 (Student’s unpaired t-test; n = 3), error bars indicate standard error of the mean.

While RI, MXE and A3SS showed some association with CAG length, especially in the striatum at 6 months of age, the SE splicing subtype showed the strongest repeat length association (see R^2^ in Supplemental_Fig_S3). Importantly, the overlap, between genes exhibiting AS alteration and expression differences at each time point and with any genotype tested, was negligible (1.2 %) (Supplemental_Fig_S4), indicating that the AS alterations were not caused by concomitant transcriptional changes or *vice versa*.

Because the striatum exhibited a more prominent alteration in aberrant AS events during the progression of the pathologic process, we then asked whether aberrant AS events were shared among a common set of striatal genes regardless of *time points* and genotype. As depicted by the circos plot in Fig. 1b, the vast majority of genes with differential AS were unique to a specific genotype and time point, though involved in redundant GO terms such as ‘synapse’, ‘cell junction’, ‘transcription’ (Supplemental_Table_S1). However, some common genes could be found among the lines with the most expanded CAG tracts (Q140 and Q175), mainly at 6 months of age (Fig. 1b). Due to the prevalence of aberrant SE events compared to other subtypes, we performed functional enrichment analysis on genes exhibiting CAG-dependent SE mis-splicing events. We analyzed the highest CAG repeat length lines (Q140, Q175) separately, while the “lower” repeat length lines (Q80, Q92, and Q111), with fewer altered isoforms, were grouped together (Fig. 1c). Functional enrichment revealed that ‘transcriptional regulation’, ‘cell junction’, and ‘synapse’ were recurrent among genotypes and the most significantly enriched terms (Fig. 1c, Supplemental_Table_S1).

Given the predominance of aberrant SE in the mouse striatum, we wanted to experimentally validate the differential exon inclusion observed using two separate lines (Q111 and Q175) in striatal samples dissected from 6 months old mice. We prioritized a list of transcripts from the AS splicing analysis involved in ‘transcriptional regulation’, ‘synapses’ processes, known pathways altered in HD, and showing a significant change in percent-spliced-in (│ΔPSI│ > 0.25) of alternative exons between Q20 controls and Q111 or Q175 (Supplemental_Table_S1). *Crem*, a cAMP-responsive element modulator (Nagamori et al. 2006), was an interesting target showing aberrant SE (ΔPSI (JCEC) = - 0.3, SE increase in *Htt* CAG expanded alleles) in both Q111 and Q175 striata. Replicate RT-PCR experiments, using an independent set of Q20, Q111 and Q175 mouse striatal samples, confirmed the altered *Crem* SE event (Fig. 1d).

### Aberrant linear alternative splicing correlates with *Htt* CAG size in murine neural progenitor cells

In order to follow how linear AS behaved during the transition from pluripotency to neuronal commitment in presence of the *Htt* CAG expansion mutation, we took advantage of a panel of mouse embryonic stem cells (mESCs) (Q20, Q50, Q92 and Q111) harboring different CAGs within a single *Htt* allele (Auerbach et al. 2001; Wheeler et al. 1999; Jacobsen et al. 2011a), closely mimicking the human HD mutations. We also included wild-type (WT) and *Htt* null alleles in which the endogenous *Htt* locus was unaltered or inactivated by the insertion of a neomycin cassette between exon 4 and 5, respectively (White et al. 1997). mESC with different genotypes were pushed toward neural differentiation following a previously reported protocol (Conti et al. 2005). Differentiated mouse neural progenitor cells (mNPC) were initially characterized by immunofluorescence and RT-qPCR, to confirm the correct reduction of pluripotency markers (i.e. *Pou5f1, Nanog*) and the induction of neural progenitor markers (i.e. *Nes, Vim*, *Msi1* and *Sox2*) (Supplemental_Fig_S5 a-b).

Following RNAseq, linear AS events for mESCs and mNPCs were analyzed by the same pipeline employed for the *in vivo* analysis, comparing the different expanded *Htt* CAG repeat lengths (Q50, Q92, and Q111) to the Q20 *Htt* CAG repeat (control). Similar to the previous results (Fig. 1a), a direct correlation between the total number of genes with aberrant AS events and CAG repeat length was observed in mNPCs but not in mESCs (Fig. 2a).

**Figure 2:**
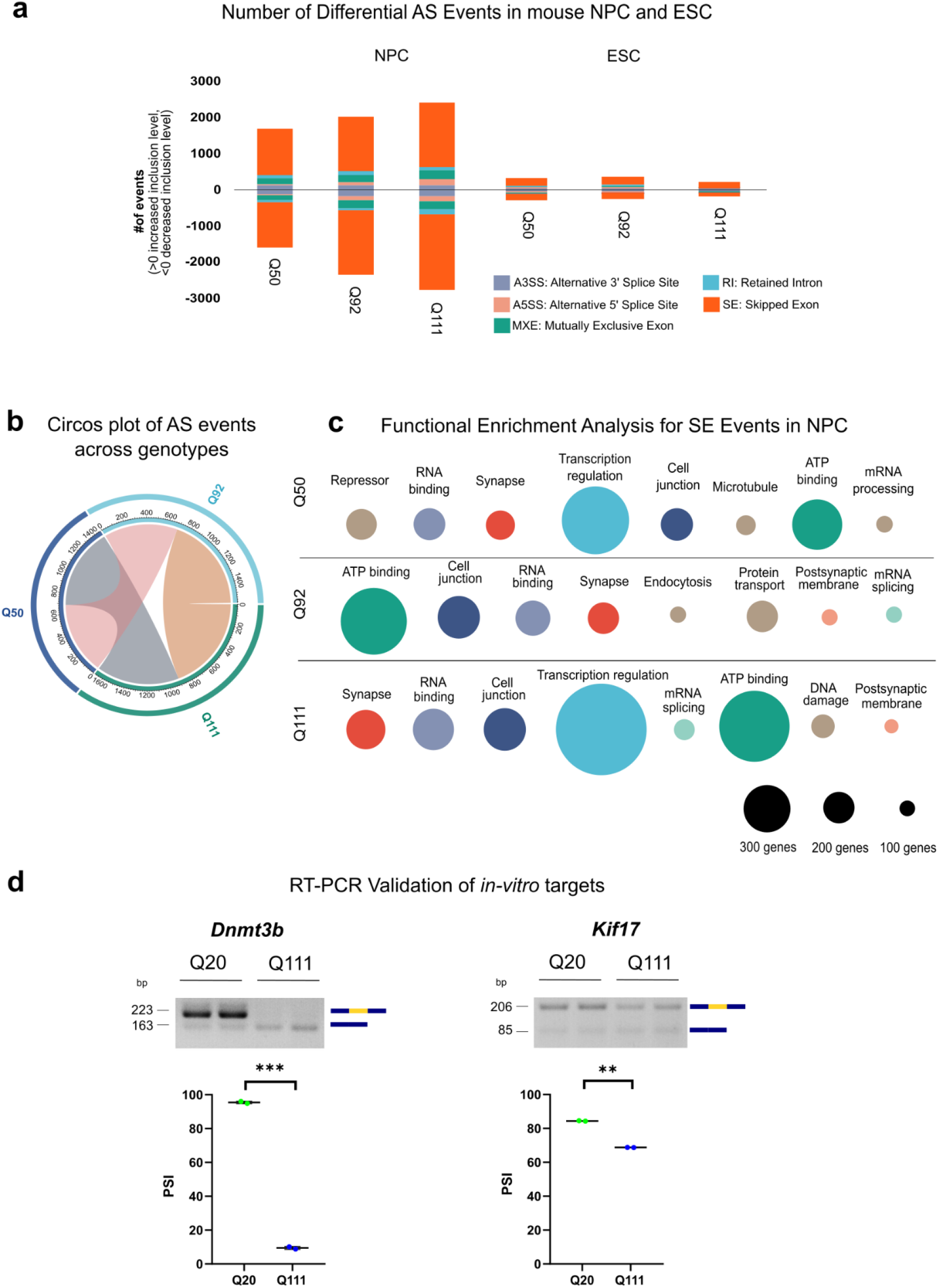
Aberrant Linear Alternative Splicing correlates with *Htt* CAG size in mNPC. **a)** The bar graph reports the number of differential AS events in the mNPC and mESC from KI models of HD, presenting 3 different *Htt*-CAG repeat lengths (Q50, Q92, Q111). The number of events is reported for each genotype and differentiation stage. The inclusion level is calculated in comparison to Q20 controls and the positive or negative values plotted in the graph. Further details in the Methods section. Each color of the bar chart represents a different AS event. **b)** Circos plot reports the number of transcripts in mNPC - presenting differential AS events - shared between different genotypes (Q80, Q92, Q111). Conditions sharing more than 50 transcripts are depicted in colors. **c)** Weighted nodes graphical representation shows the functional enrichment analysis for transcripts presenting significant skipped exon (SE) events in mNPC. Each genotype is shown separately. Nodes’ size legend is depicted at the bottom. Nodes are ordered by adjusted p-values using Benjamini Hochberg (<0.05). The same enrichment terms shared between genotypes are presented with nodes of the same color, while unique terms are colored with light brown. **d)** Representative agarose gel images (top) and dot-plots (bottom) report the RT-PCR results of *in vitro* AS validation for *Dnmt3b* and *Kif17*, selected transcript targets. RT-PCR assay and quantification were performed on RNA from Q20 and Q111 mNPCs. *Dnmt3b* and *Kif17* isoforms presenting inclusion or exclusion of the variable exons (in yellow) are visualized and quantified. Dot plots report the PSI, percent-spliced-in. **P < 0.01, ***P < 0.001 (Student’s unpaired t-test; n = 3), error bars indicate standard error of the mean.

Interestingly, similar results were obtained from the analysis of AS events in *Htt* null mNPCs but the affected genes were mostly different (only 11% were overlapping) and unique to KO or KI condition (Supplemental_Fig_S6). Similar to the *in vivo* models, the highest proportion of aberrant AS events was the SE subtype and increasing *Htt* CAG size was more strongly linked to decreased rather than increased inclusion levels (Fig. 2a). Importantly, also in the *in vitro* models, the AS alterations were vastly not associated with concomitant transcriptional changes, with minor overlaps (20.8%) detected between differentially expressed (DE) and AS-affected genes among genotypes and differentiation time points (Supplemental_Table_S2). Consistent with a more homogeneous cellular population, mNPCs showed a clear overlap between genes showing/exhibiting aberrant splicing (any subtype) across different genotypes (Fig. 2b). Moreover, functional enrichment of genes showing/exhibiting mis-splicing events showed that ‘RNA-binding’, ‘mRNA processing’, ‘splicing’, ‘kinase’ and ‘synapse’ were the most significantly enriched terms (Supplemental_Table_S2). Due to the prevalence of SE subtype among the mis-spliced events, a dedicated functional enrichment analysis was performed for each genotype separately. The analysis highlighted terms such as ‘transcriptional regulation’, ‘ATP binding’, ‘RNA binding’ and ‘synapse’ as recurrent among the genotypes (Fig. 2d, Supplemental_Table_S2) and mirroring the *in vivo* data (Fig. 1c). Similarly, we prioritized transcripts involved in the indicated biological pathways with statistically significant SE events with │ΔPSI │ > 0.5 between Q20 and Q111 NPC for RT-PCR experimental validation. *Dnmt3b* (SE (Q111_JCEC) ΔPSI = −0.86, SE increase in *Htt* CAG expanded alleles), involved in maintenance of DNA methylation and transcriptional regulation processes (Okano et al. 1999) and *Kif17* (Kinesin Family Member 17) (SE (Q111_JCEC) ΔPSI = −1.0, SE increase in *Htt* CAG expanded alleles) with ATPase and microtubule motor activities (Wong-Riley and Besharse 2012) were prioritized. Replicate RT-PCR experiments validated the altered SE events (Fig. 2d).

### *Htt* CAG size dictates a specific linear AS signature in neural cells

Because of the similarities in the AS results between the *in vivo* and *in vitro* HD models, we sought to directly intersect the genes exhibiting SE in the mouse striatum and mNPCs. While the majority of SE events in the striatum were age-specific (Fig. 3a), some more overlap was observed at 6 and 10 months of age (123 genes shared, 13% of total). Strikingly, however, *Htt* CAG expansion caused a specific AS disruption in genes common in the *in vivo* and *in vitro* HD models (45% of the total striatal SE events, p-value: 1.16^-163, Fisher’s exact test) (Fig. 3b), which were mainly involved in ‘cell junction’ and ‘synapse’ enriched GO terms (Supplemental_Table_S1-2).

**Figure 3.**
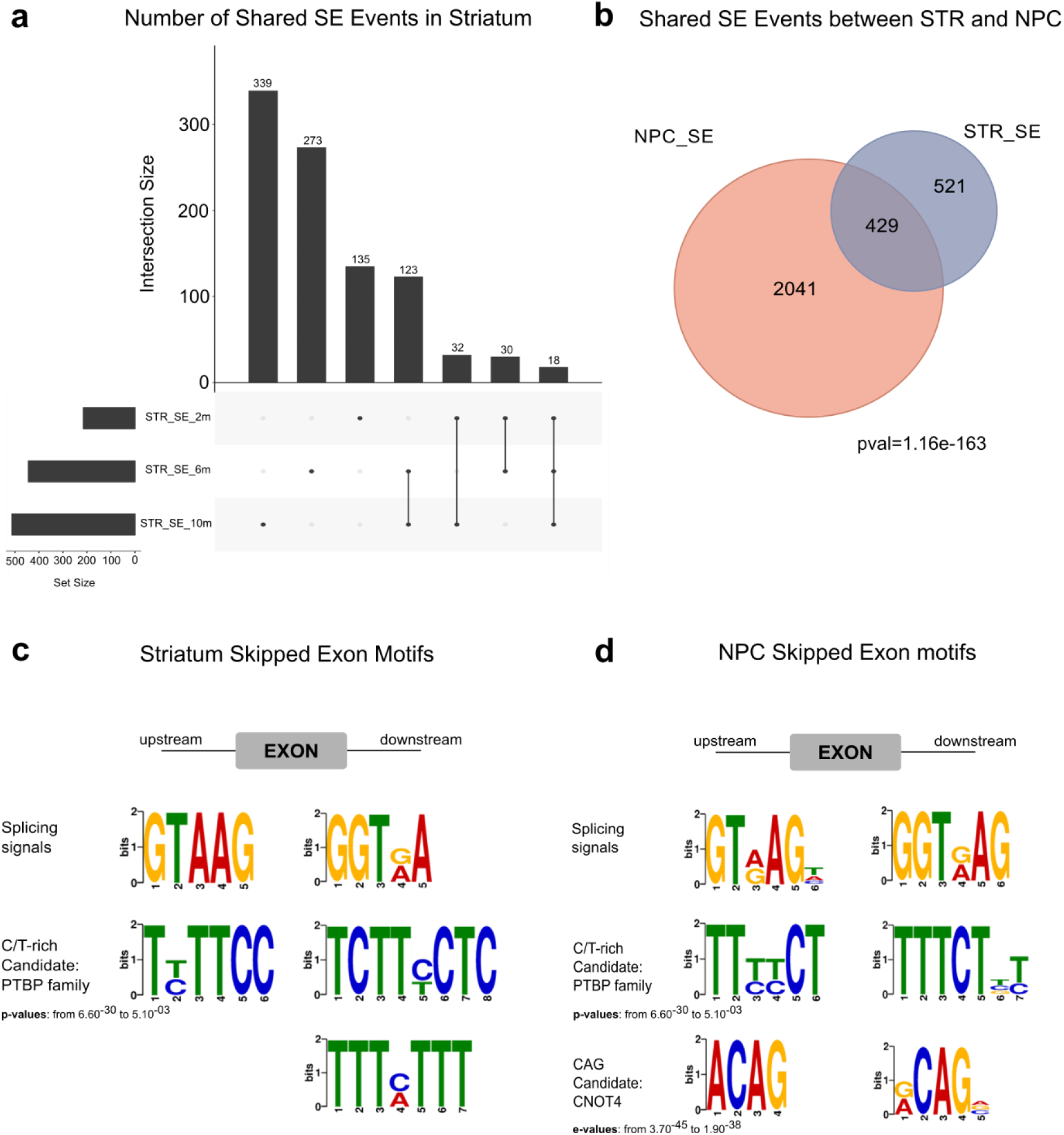
*Htt*-CAG size dictates a specific linear AS signature in neural cells. **a)** The upset plot displays the number of SE events shared among different time points (2, 6 and 10 months) within the mouse striatum. The number of events within each intersection is presented in the vertical bars. Intersection groups (lines) or single time points (dots) are shown in the lower panel. Sample set size is indicated at the bottom left of the panel. **b)** The Venn diagram reports the comparison between striatal and mNPC SE events. All genotypes (Q80, Q92, Q111, Q140 and Q175) and time points (2, 6 and 10 months) for striatal districts and all genotypes (Q50, Q92, Q111) for mNPC were combined. Shared SE events (45% of the total striatal SE events, p-value: 1.16^-163, Fisher’s exact test) are indicated in the intersection. **c-d)** Motif analysis identifies the splicing factors and/or RBP binding sites in the ± 100bp upstream and downstream adjacent regions to the alternatively spliced exons for the striatum (c) and for mNPC (d). The P-value of enrichment testing for individual motifs in each data set is indicated. The candidate binding splicing factor and/or RBP family is shown.

Importantly, by inspecting the regions adjacent to the differentially skipped exons (+/- 100bp) in both mouse striatum and mNPCs, we detected a specific and significant enrichment for C/T rich motifs (p-values from 6.60^-30 to 5.10^-03), typically bound by polypyrimidine tract binding proteins (PTBP) (Ray et al. 2013) (Fig. 3c, d and Supplemental_Table_S3). Furthermore, a shorter, CAG-rich motif was found (p-values from 3.70^-45 to 1.90^-38), which could possibly be bound by CNOT4 (Lau et al. 2009).

### *Htt* CAG mutation impacts circRNA biogenesis in mouse neural progenitors

Considering that AS is fundamental to the biogenesis of circRNAs, we asked whether the *Htt*-CAG expansion could be also associated with defective back-splicing in the *in vitro* system. First, the analysis of the percentage of circRNAs originating from each chromosome generally complied with the expected ratio relative to transcript mass in both mESCs and mNPCs (Supplemental_Fig_S7 a-b). CircRNA reads (average reads/sample = 3825) were then quantified in all samples by means of the DCC platform (Cheng et al. 2016) (Supplemental_Table_S4-5). As shown in Fig. 4a, we observed, for all genotypes, a higher number of expressed circRNAs in mNPC than in mESC samples with a corresponding higher circRNA fraction of total transcript mass (circRNA reads/circRNA + host transcript reads, Supplemental_Fig_S7c). This trend was specific for circRNAs, given that when evaluating small RNAs [miRNAs, mitochondrial transfer RNAs (Mt-tRNAs), processed pseudogenes, ribosomal RNAs (rRNAs), small nucleolar RNAs (snRNAs), small Cajal body-specific RNAs (scaRNAs), To be Experimentally Confirmed (TEC), transcribed processed pseudogenes] (average aligned reads/sample = 1.04M, see methods, Supplemental_Table_4-5), we observed a general reduction in mNPCs compared to mESCs (Fig. 4b). Specifically, miRNAs and processed pseudogenes subtypes appeared to be strongly affected by neural differentiation, considerably reduced in mNPCs (Fig. 4b and Supplemental_Fig_S8). While more than half of circRNA species derived from coding sequences (CDSs) and 5’-untranslated regions (5’-UTRs) in both mESCs and in mNPCs (Fig. 4c), their spliced lengths were significantly longer in mNPCs (Fig. 4d).

**Figure 4.**
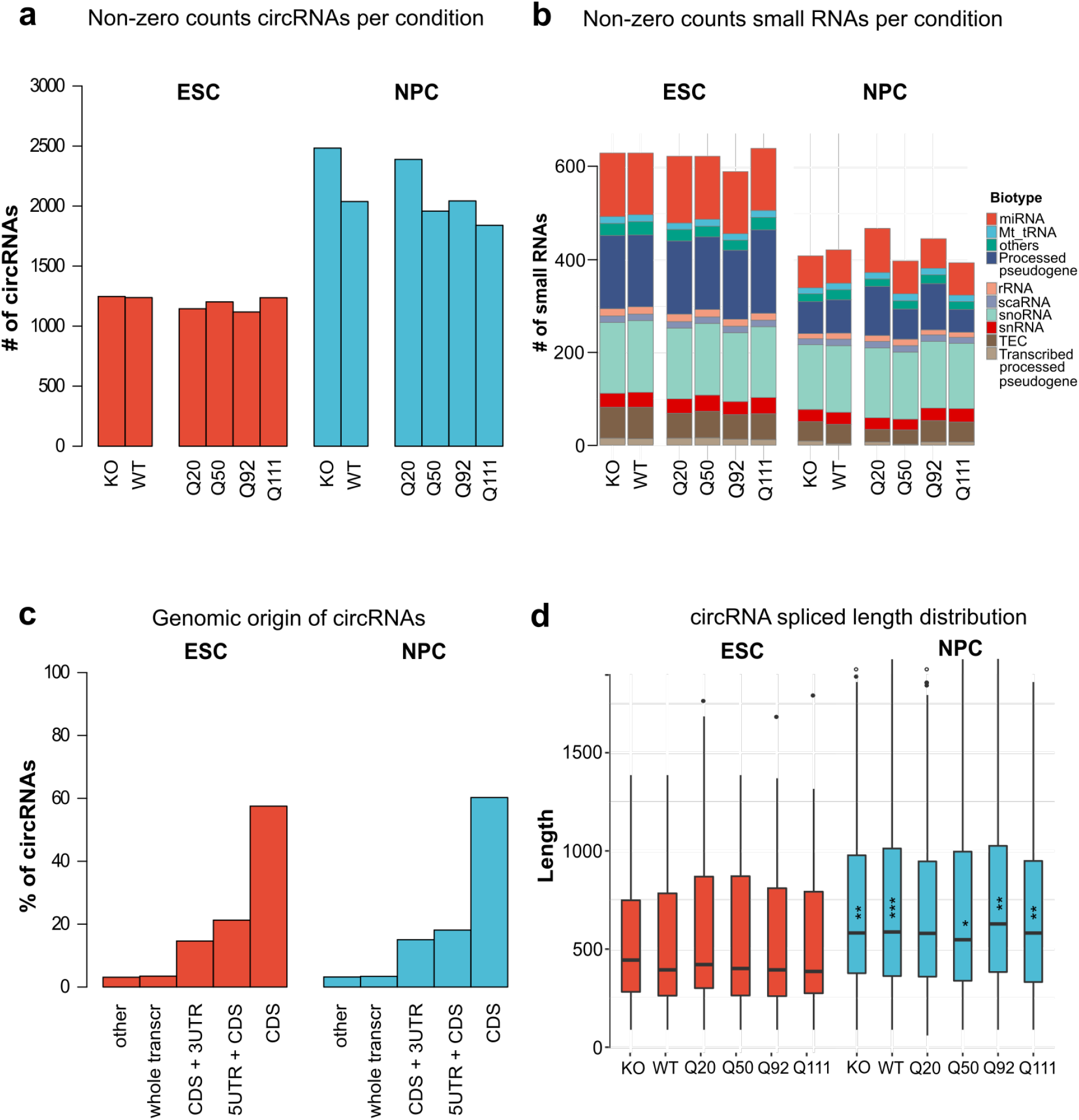
Neural differentiation impacts on circRNAs and small-RNAs biogenesis. **a)** The bar chart shows the number of detected circRNAs (>1 count/sample, see also Methods), comparing pluripotent (mESC) and neural committed progenitors (mNPC). Different *Htt* genotypes are presented: *Htt* double knock-out (KO) and a series of 4 *Htt*-CAG expansion alleles (Q20, Q50, Q92 and Q111). Average circRNA count from two biological replicate experiments is plotted. **b)** The color bar graphs report the number of small RNAs (>1 count/sample, see also Methods) comparing mESC and mNPC conditions. *Htt* genotypes as in a). Different classes of small RNAs are examined. Abbreviations as follows: microRNAs (miRNAs), Mitochondrial transfer RNAs (Mt-tRNAs), processed pseudogenes, ribosomal RNAs (rRNAs), small nucleolar RNAs (snRNAs), small Cajal body-specific RNAs (scaRNAs), To be Experimentally Confirmed (TEC), transcribed processed pseudogenes. **c)** The bar chart reports the percentage of circRNAs derived from specific transcripts areas: transcript coding sequence (CDS), 3’ or 5’ untranslated regions (3UTR/5UTR), whole transcript or other. The data compare mESC and mNPC conditions. All *Htt* genotypes were combined. **d)** The box plot displays the spliced length distribution [average ± standard deviation (SD)] for circRNAs in the various conditions of mESC and mNPC cells. *Htt* genotypes as in a). Wilcoxon test p-values of the difference between corresponding conditions of mESC and mNPC are shown as stars (* < 0.05, ** < 0.01, *** < 0.001).

We then analyzed the changes in circRNA expression between samples of increasing CAG length. As shown in Fig. 5a, while in mESC the number of increasing and decreasing circRNAs was almost equal, circRNA expression in mNPCs predominantly decreased in Q111 vs Q20 samples (92 increasing vs 478 decreasing). Genes harboring differentially expressed circRNAs were over-represented in pathways and functions such as ‘cGMP-PKG signaling pathway’, ‘Lysosome’ and ‘DNA binding’ (Supplemental_Table_S4). Interestingly, the complete absence of huntingtin in mNPCs elicited an opposite effect on circRNA production with the majority of circular RNA molecules turned on in *Htt* KO mNPCs (Supplemental_Fig_S9 a-b).

**Figure 5.**
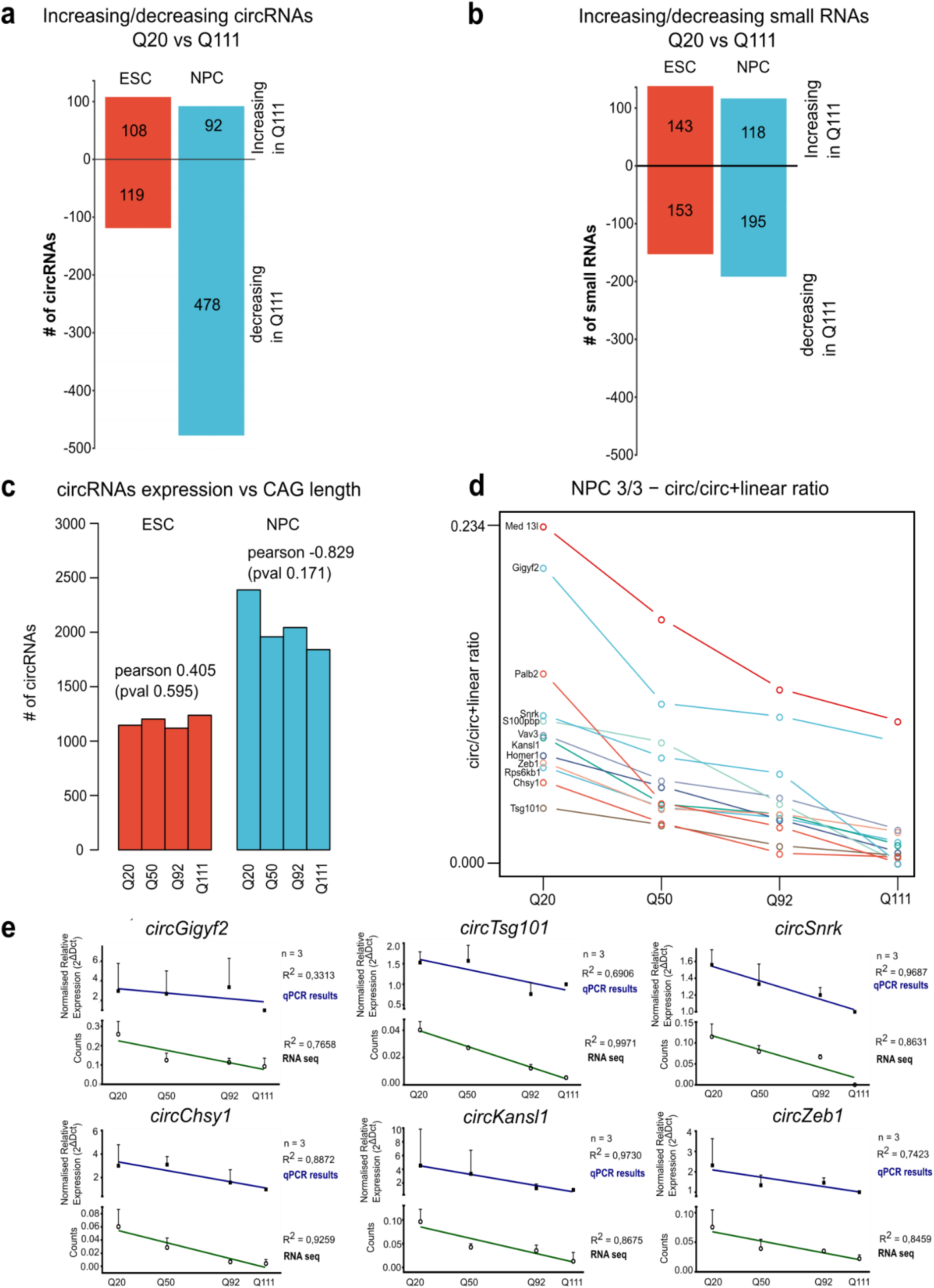
*Htt*-CAG dependent reduction of circRNAs in mNPC. **a)** The bar graph reports the number of circRNAs differentially expressed between *Htt* Q111 versus Q20 genotypes. The comparison is presented for pluripotent (mESC) and neural committed progenitors (mNPC). The number of circRNAs increasing (*Increasing in Q111* - upper part of the plot), and decreasing their expression in Q111 versus Q20 (*Decreasing in Q111* - lower part of the plot) is depicted. **b)** The bar chart shows the number of small-RNAs differentially expressed between *Htt* Q111 versus Q20 genotypes. The comparison is presented for pluripotent (mESC) and neural committed progenitors (mNPC). The number of small-RNAs increasing (*Increasing in Q111* - upper part of the plot), and decreasing their expression in Q111 versus Q20 (*Decreasing in Q111* - lower part of the plot) is depicted. **c)** The bar chart reports the total number of circRNAs expressed at each differentiation stage (mESC and mNPC) and for each *Htt*-CAG genotypes (Q20, Q50, Q92 and Q111). A negative correlation between the number of expressed circRNAs and *Htt-*CAG length is observed (although not nominally significant) in mNPC, but not in mESC cells. Pearson’s correlation R = −0829, p-value = 0.171. d) Line plot describes the expression pattern of the 12 circRNAs of mNPC selected by three criteria (decreasing expression and negative correlation with *Htt-* CAG, significantly different expression by circTest-see Methods for further details). *Htt*-CAG expansion alleles (Q20, Q50, Q92 and Q111) are presented. **e)** The line charts report the ratio (circ/linear) of the expression levels for the selected circRNAs candidates in mNPC across different *Htt*-CAG genotypes. RT-qPCRs results of 3 biological replicates are plotted as normalized circ/linear expression (2^^ΔΔCT^) and average ± standard deviation (SD) is presented. *Htt*-Q111 genotype – with lower circ/linear ratios - was used as relativizing condition. The linear trend lines are presented for the qPCR validation (blue line) and the corresponding RNAseq data (green line) (qPCR, filled symbols; RNAseq, empty symbols). RNAseq data are presented as normalized transcript counts (counts). Pearson’s correlation R square values are indicated for the two regressions (qPCR and RNAseq).

To uncover a possible link between *Htt* CAG expansion and decreased circRNA expression, we counted the number of expressed circRNAs in each *Htt* CAG genotype. Although not nominally statistically significant (p = 0.171), possibly because of the reduced number of genotypes examined (n = 4), a high inverse correlation (Pearson’s coefficient = −0.829) was observed in mNPCs (Fig. 5c and Supplemental_Fig_S7c). Such a trend was absent in mESCs. We then selected a stringently defined subset of differentially expressed circRNAs whose expression was (i) monotonically decreased with increasing CAG lengths; (ii) significantly (p < 0.05) changed according to Pearson’s correlation coefficient and (iii) significantly different as defined by CircTest algorithm (Cheng et al. 2016). 12 circRNAs fulfilling these criteria and showing a negative trend of expression with increasing *Htt* CAG length (Fig. 5d), were used as circRNA candidates for experimental RT-qPCR validation. Biological triplicates of ribo-depleted RNAs for the 4 *Htt* CAG repeat genotypes were used for quantification of linear and circular isoforms (circRNA/circRNA+linear ratio). We experimentally confirmed the expected expression changes, i.e. circRNA expression linearly decreasing from Q20 to Q111, in 6 of 10 targets (Fig. 5e). Significantly, circ*Snrk* and circ*Kansl1,* previously characterized for their function in regulating apoptosis (Zhu et al. 2021) and as miRNA sponges (Wang et al. 2020), showed the highest CAG correlation Pearson’s R^2^ and most significant p-values (circ*Snrk* R^2^ = 0.9687, p = 0.0313 and circ *Kansl1* R^2^ = 0.9730, p = 0.0270) (Fig. 5e).

### Linear and back-splicing alterations correlate with direct and indirect mis-regulation of RNA-binding proteins and splicing factors

In order to deepen our understanding of the effects of *Htt* CAG expansion on linear and back-splicing regulation and to gather new mechanistic insights, we focused on miRNAs, small RNAs subtype, which are very sensitive to neural differentiation (Fig. 4b) and [already] reported to control the stability of RNA-binding proteins and splicing factors in neurons (Weiss et al. 2015; Fukao et al. 2015; Gardiner et al. 2015). Although miRNA expression strongly decreased following neural differentiation (Fig. 4b and Supplemental_Fig_S8), we asked whether we could identify differentially expressed miRNAs affected by the expression of *Htt* CAG expansion in mNPC. Thus, we compared Q20 versus Q111 mNPCs and identified 9 significantly differentially expressed miRNAs (Supplemental_Table_S5). By exploiting the mirDB database (Chen and Wang 2020), we inferred their mRNA targets and intersected them with mouse splicing factors and RNA-binding proteins (RBPs) lists (see methods and Suppl. Table 5). We identified 73 transcripts (Fig. 6a), known RNA-binding proteins and splicing factors (of the *Celf, Elav, HnRNPs, Mbnl, Ptbp, Srsf* families). These were also targets of miRNAs dysregulated by mutant huntingtin and showing expression levels changes with │Log2FC│ > 0.5 and a significant p-value (p < 0.05) (Fig. 6a, Table). Importantly, significant overlap and enrichment was also identified between miRNA targets and transcripts showing differential alternative splicing or back-splicing in mNPC (Supplemental_Fig_S10 and Supplemental_Table_S7), suggesting a possible role for miRNAs in the regulation of AS and back-splicing events. Strikingly, *Ptbp3*, usually expressed in thymus, lymph nodes and digestive system (Uhlén et al. 2015) whose binding motifs were significantly enriched in the ± 100bp adjacent to the differentially skipped exons in both the *in vivo* and *in vitro* systems (Fig. 3c,d), was significantly transcriptionally upregulated (Fig. 6a, Table).

**Figure 6.**
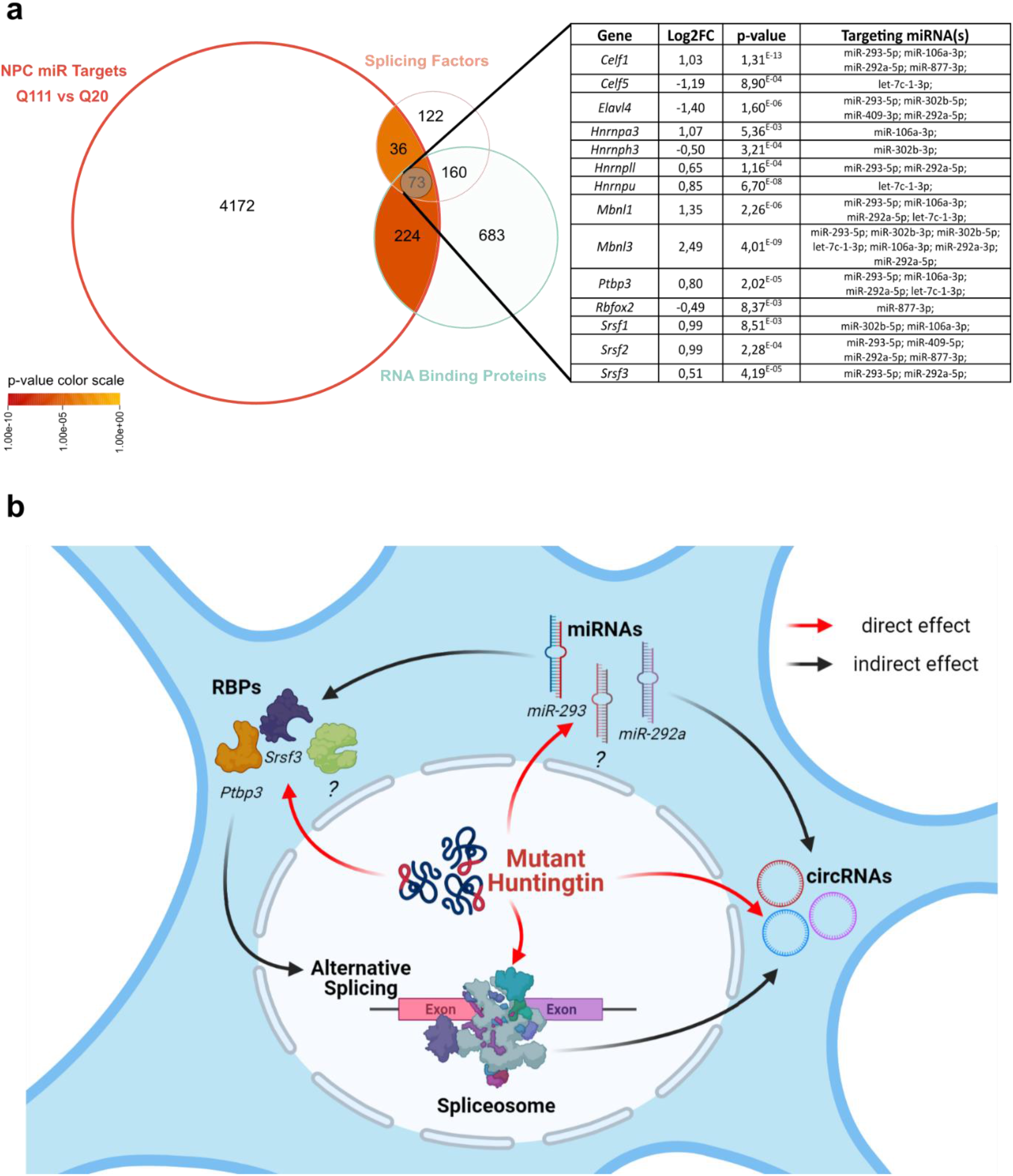
Linear and back-splicing alterations correlate with direct and indirect mis-regulation of RNA-binding proteins and splicing factors. **a)** The Venn diagrams reports the overlap between mutant huntingtin’s dysregulated miRNA targets (4505 transcripts targets of 9 dysregulated miRNAs comparing Q20 versus Q111 *Htt*-CAG genotypes in mNPC, see Methods and Supplemental_Table_S5), the list of splicing factors (391) and RNA binding proteins (1140) (see Methods and Supplemental_Table_S5). The enrichment scores observed for the different lists of splicing regulators is represented as color-coded within the intersection areas. Color-coded scale bar on the bottom left corner of the figure. A subset of transcripts with splicing regulator or RBP functions which are targets of mutant huntingtin’s dysregulated miRNAs (smaller intersection, # 73) are highlighted in the left table. Log fold change (LogFC) transcripts difference in the reference conditions (mNPC Q20 versus Q111), associated p-value and targeting miRNAs are shown. **b)** The scheme summarizes the main findings and the proposed mechanistic hypotheses of the current study, supporting a complex, direct and/or indirect, mode of mutant huntingtin regulation of canonical linear and circular RNA-producing splicing. The summary scheme was created by using BioRender.com.

Taken together, while we cannot exclude direct transcriptional alterations of RBPs and splicing factors by mutant huntingtin, we identified a pool of dysregulated miRNAs, sensitive to the *Htt* mutation and possibly targeting linear and back-splicing factors. Thus, our data support a complex direct and indirect mode of mutant huntingtin regulation of canonical linear and circular RNA-producing splicing (Fig. 6b).

## DISCUSSION

Alterations in the choice of splice sites resulting in proteins with different structures and functions, altered mRNA localization, translation or decay is crucial for the complexity of the mammalian nervous system (Li et al. 2007; Zheng and Black 2013; Raj and Blencowe 2015; Darnell 2013). Splicing defects impacting the functionality of mature neurons are increasingly implicated in neurological and neurodegenerative diseases. Thus, there is an increasing need to better understand these regulatory processes.

Since the discovery of the causal genetic mutation underlying HD, great efforts have been made to uncover the functions of both wild-type and mutant huntingtin, which is now thought to act in a truly pleiotropic manner. The challenge nowadays is to reveal which of these pathways might exert a crucial early role in the onset and progression of HD pathology.

Here, we investigated how RNA processing and, specifically, alternative splicing is affected by mutant huntingtin. By resourcing to publicly available RNA-seq data from *in vivo* HD KI mice (Q20, 80, 92, 111, 140 and 175) with different CAG lengths (Langfelder et al. 2016) and our newly generated RNA-seq from an isogenic panel of mouse ESCs and NPCs, we uncovered a neural specific, CAG-length dependent alteration in alternative splicing. Exon skipping and, to a lesser extent, intron retention events were primarily altered. Notably, the increased total number of AS events was significantly correlated with CAG length (and to a lesser extent with age) in the striatum, but not within the cortex and liver, hence supporting preferential striatal vulnerability. Interestingly, significant changes in the AS pattern could already be detected prior to overt behavioral phenotypes stages (2 months). While previous reports correlated mutant huntingtin expression to mis-splicing events, locally affecting the *Htt* locus (Neueder et al. 2017; Sathasivam et al. 2013; Schilling et al. 2019), and also more generally altering the whole brain’s transcriptome (Lin et al. 2016; Elorza et al. 2021), this is the first evidence of a direct correlation between the number of *Htt* CAG repeats and the degree of AS, extending the repertoire of molecular phenotypes directly linked to the CAG expansion. The concept of CAG-dependency is currently extensively investigated to gain insights into how *Htt* CAG repeat length might modify HD pathogenesis (Langfelder et al. 2016; Galkina et al. 2014; Reis et al. 2011; Seong et al. 2005). Notably, our findings supporting a primary CAG length-dependent effect in striatum – with less involvement of cortex [and liver] – is well in line with previous observations that highlighted CAG length-dependent modules of co-expressed genes (Langfelder et al. 2016).

Similarly, the results obtained from the AS analyses, on the isogenic *in vitro* KI (Auerbach et al. 2001; Wheeler et al. 1999; White et al. 1997; Duyao et al. 1995; Jacobsen et al. 2011b) system in the transition from pluripotency to neural committed progenitors, provided an independent, genome-wide validation of the AS *Htt* CAG correlation. Further, we confirmed an increase of AS and SE events with increasing CAG length. However, this phenotype was limited to mNPCs, which also exhibited a greater number of total AS events relative to mESCs. These observations globally suggest that (i) neural progenitors activate prominent neural splicing processes (Weyn-Vanhentenryck et al. 2018), and (ii) the correlation with CAG length likely requires factors expressed in the neural lineage. Functional enrichment analyses for transcripts experiencing SE events, *in vivo* and *in vitro*, revealed GO and pathways already associated with HD (Labbadia and Morimoto 2013; Hodges et al. 2006; Kuhn et al. 2007), with prevalence of transcriptional and neural-related functions. Taken together, these findings suggest that linear splicing becomes progressively dysregulated as CAG length increases, specifically in neuronally-committed cells and adult striatum, thus possibly contributing to HD pathogenesis.

It has been ascertained that AS regulation is also crucial to the biogenesis of circRNAs, stable, neuronally-enriched, circular non-coding RNAs (Ashwal-Fluss et al. 2014; Ivanov et al. 2015), with important roles during neural development (You et al. 2015; Venø et al. 2015a) and in brain function (Piwecka et al. 2017; Lu et al. 2019). While their contribution to brain pathologies and specifically to neurodegenerative diseases is becoming recognized (Jia et al. 2020; Zhang et al. 2020; Dube et al. 2019), circRNAs remain largely neglected in HD. We aimed to provide a first characterization of back-splicing regulation in the presence of mutant huntingtin expression, exploiting the isogenic *in vitro* KI system. In contrast to what was observed for linear splicing, we detected a general decrease in circRNA abundance with increasing CAG length. While the impact of mutant huntingtin on circRNAs biogenesis is remarkable when comparing the two extreme genotypes (Q20 versus Q111), the genes affected by AS and back-splicing are largely different, and also enriched in different GO and pathways terms. However, once again, the more striking phenotype appeared to be confined to neural progenitors with minimal changes in mESC.s It is unclear at present whether these circRNAs play known or novel regulatory roles (Wilusz 2018), and whether their depletion might contribute to HD pathogenesis.

Taken together, our observations demonstrate an opposite correlation of linear AS and back-splicing with the number of CAG repeats, thus suggesting a possible link between the two splicing types, with contrary/reversed regulation. Previous studies reported exon skipping as a promoter of skipped-exon circularization (Kelly et al. 2015). However, other data suggested that alternative splicing is likely in competition with back-splicing (Holdt et al. 2018; Ashwal-Fluss et al. 2014). Our results support the latter idea, where a global increase of SE events downregulates back-splicing efficiency although it remains unclear whether the *Htt* CAG mutation primarily affect linear splicing, exon circularization, or both simultaneously. The effect exerted by the *Htt* CAG expansion on circRNA biogenesis in mNPCs, might be exacerbated by the higher number of circRNAs detected at this developmental stage, already reported in a number of studies (Izuogu et al. 2018; Szabo et al. 2015; Venø et al. 2015b). On the other hand, it might suggest that the *Htt* CAG expansion could, directly or indirectly, regulate some neural splicing factors also implicated in back-splicing control. the *Htt* CAG expansion, thus, might induce an abnormal interaction and/or alter the expression levels of small RNAs, splicing factors or RBP, specifically in neural-lineages (Murthy et al. 2019), in turn affecting the two types of splicing.

Here we explored a network of miRNAs and their targets to expose some mechanistic insights. miRNAs revealed to be altered by neural differentiation and, to a lesser extent, by the *Htt* CAG expansion. Interestingly, a fraction of the targets of mutant huntingtin’s-dysregulated miRNAs, were significantly enriched for RBP and splicing factors, with members of the PTBP, CELF, SRSF and hnRNPs families, also showing transcriptional alterations (Fig. 6a, Table). This suggests that these AS regulatory elements might be indirectly (via miRNAs alteration) or directly dysregulated by *Htt* CAG expansion (Fig. 6 b). Importantly, targets of mutant huntingtin’s-dysregulated miRNAs also revealed a significant enrichment for transcripts exhibiting SE and back-splicing events in mNPCs, supporting the existence of a complex RNA regulatory network by mutant huntingtin (Fig. 6 b).

In conclusion, we identified splicing and back-splicing as highly impacted molecular alterations in HD model systems. Although previous reports associated mutant huntingtin expression to linear splicing dysregulation (Lin et al. 2016; Schilling et al. 2019; Elorza et al. 2021), this is the first study reporting the relationship between magnitude of AS alterations and *Htt* CAG length and the very first evidence of circRNA deregulation in HD. Moreover, the identification of aberrantly regulated miRNAs, RBPs, and splicing factors, specifically in neural committed cells and in combination with *Htt* mutations, proposes novel molecular players contributing to HD pathogenesis and delineating new targets of therapeutic intervention.

## METHODS

### Generation and characterization of mNPC with different *Htt* CAG sizes

The isogenic panel of wild-type, *Htt* null *Hdh*^ex4/5/ex4/5^ and heterozygous *Htt* CAG knock-in *Hdh*^Q20/7^, *Hdh*^Q50/7^, *Hdh*^Q92/7^ and *Hdh*^Q111/7^ mESCs, kindly provided by Dr. Marcy E. MacDonald (Massachusetts General Hospital and Harvard Medical School, Boston, USA), were cultured as described previously (Auerbach et al. 2001; Wheeler et al. 1999; White et al. 1997; Duyao et al. 1995; Jacobsen et al. 2011b). Pluripotent cells were maintained in KnockOut DMEM (Gibco), supplemented with 15% of ESC-grade FBS (Gibco), 2 mM L-glutamine (Gibco), 100 U/ml Penicillin/Streptomycin (Gibco), 1% Non-essential Amino Acids (Gibco), 0.1 mM 2-mercaptoethanol (Sigma) and 1000 U/ml of leukemia inhibitory factor (LIF) (Voden), on plates coated with 0.1% gelatin (Millipore) or on a feeder layer of CF-1 IRR mouse embryonic fibroblast (TebuBio).

Neural differentiation was performed as previously described (Conti et al. 2005). Briefly, self-renewing mESCs were dissociated and plated onto 0.1% gelatin-coated plates at a density of 0.5–1.5 10^4^cells/cm^2^ in N2-B27 medium (Ying et al., 2003). After 7 days, cells were detached using Accutase (Thermo Fisher Scientific) and plated on 3 µg/ml laminin-coated dishes in neural stem (NS) expansion medium [composed by Euromed-N (Euroclone) supplemented with 20ng/ml FGF (R&D) and EGF (Sigma), 1% N-2 Supplement (Gibco), 2 mM L-glutamine (Gibco) and 100 U/ml Penicillin/Streptomycin (Gibco)] (Conti et al. 2005). Mouse neural progenitor cells were routinely passaged 1:2-1:4 every 3-5 days using Accutase and maintained in NS expansion medium on laminin-coated plates (Sigma, 3 μg/ml). Both mESCs and mNPCs were incubated at 37 °C and 5% CO2. The pluripotency of mESCs and their differentiation to mNPCs was evaluated by RT-qPCR using stage specific markers (full list of primers used in Supplemental_Fig_S6).

### Immunofluorescence

Cells were seeded on a 24-well the day before the experiment and then fixed with 4% PFA for 15 minutes at room temperature, permeabilized with 0.5% Triton X-100 in 1X PBS and incubated with blocking solution (0.3% Triton X-100, 5% FBS in 1X PBS) for 1 hour. Primary antibody incubation was carried out at 4 °C overnight, followed by three washes with 1X PBS. Proper secondary antibodies were eventually used and images acquired using a confocal microscope (Leica TCS SP5). The following antibodies were used: rat anti-Nestin (Santa Cruz Biotechnology), rabbit anti-Sox2 (GeneTex), goat Alexa Fluor 546 and 647 (Life Technologies). Nuclei were stained using Hoechst 33342 (Life Technologies), diluted 1:20000 in 1X PBS.

### Tissues dissection and isolation

All animal experiments were conducted to minimize pain and discomfort, under approved Institutional Animal Care and Use Committee (IACUC) protocol of the Italian Ministry of Health (project authorization n. 781/2016-PR) and the Massachusetts General Hospital. 3 *Htt* KI mouse lines with an *Htt* CAG repeat knock-in allele (*Htt*^Q20^, *Htt*^Q111^ and zQ175) (C57BL/6 J inbred background) have been described previously (Wheeler et al. 1999; Lee et al. 2011; Grima et al. 2017; Menalled et al. 2003). Mice were maintained as heterozygotes and genotyped according to previously published protocols (Wheeler et al. 1999; Lee et al. 2011; Grima et al. 2017; Menalled et al. 2003).

Mice were sacrificed by CO_2_ asphyxiation followed by cervical dislocation. The brain regions of interest (striatum and cortex) were dissected on ice, rapidly removed, snap frozen and stored at −80 °C for further use. Each group/genotype included 3 males and 3 females to avoid sex-dependent bias, were collected at 6 months of age.

### RNA extraction and reverse transcription

Isolation of total RNA from tissues and cells was performed using TRIzol-based extraction method according to the manufacturer’s protocol (Thermo Scientific). RNA was resuspended in nuclease-free water and quantified by a Nanodrop 2000 spectrophotometer (Thermo Scientific). The quality of RNA was estimated by using the RNA 6000 Nano or Pico Bioanalyzer 2100 Assay (Agilent). RNA samples with RNA integrity number between 6.8 and 9.5 were used. Unless otherwise noted, 1 µg of total RNA was reverse transcribed using SensiFAST cDNA Synthesis Kit following the manufacturer’s protocol (Bioline).

### Primer design

Transcripts were searched on Ensembl genome browser 95 (https://www.ensembl.org/Mus_musculus/Info/Index) and primers were designed on the coding sequence of each gene. Primers for end-point PCR were generated using the default setting on Primer3web (http://primer3.ut.ee/) and those for quantitative PCR by using Roche Universal ProbeLibrary for Mouse (https://lifescience.roche.com/). The list of primers used in this study is presented in Supplemental_Table_S6.

### Quantitative PCR analysis

Quantitative polymerase chain reaction (qPCR) was performed using SensiFAST™ SYBR No-ROX Kit (Bioline, BIO-98020) according to the manufacturer’s instructions. Data was normalized with RNA Polymerase II subunit A (*Polr2a*) as housekeeping gene (ΔCT) and analyzed with the 40-ΔCT method. The 40-ΔCT method was used to distinguish between down-regulation and overexpression of markers of mESCs and mNPCs

### Validation of linear AS candidates’ transcripts

Candidate transcripts presenting differential linear AS among the different conditions were filtered on the basis of two parameters: (i) the percentage difference between the length of the target exon – included or excluded - and the invariant adjacent exons should be higher than 25% and (ii) the percentage of target exon inclusion/exclusion difference between *Htt*^Q111^ and *Htt*^Q20^ or *Htt*^Q175^ and *Htt*^Q20^ mice should be higher than 25%. For the selected candidates, primers were designed targeting the invariant adjacent exons to amplify the two splicing isoforms. RT-PCR was then performed using Phusion Green Hot Start II High-Fidelity PCR Master Mix (Thermo Scientific) and 20-100 ng cDNA. PCR amplicons were resolved on 2% agarose gels pre-stained with Xpert Green DNA Stain (Grisp) in 1× TBE buffer (89 mM Tris-borate and 2 mM EDTA, pH 8.3). The intensity of the bands was quantified using ImageJ software (NIH, Bethesda, MD) and the percent spliced-in value was calculated as (intensity of band with exon inclusion divided by the sum of the intensity of bands with both exon inclusion and exclusion) x 100. Replicate experiments (2 or 3) were carried out and statistical analysis was conducted using Student’s unpaired t-test. P ≤ 0.05 (*) or P ≤ 0.01 (**)

### Validation of expression changes for selected circular RNA candidates

For circular RNA validation experiments, 10 μg of total RNA was depleted of rRNAs by using the Human/Mouse RiboMinus Transcriptome Isolation Kit (Invitrogen). Ribosomal RNA depletion was confirmed by an Agilent 2100 Bioanalyzer (Agilent). Ribo-depleted RNA was reverse-transcribed and quantitative PCR was performed by employing iTaq Universal SYBR® Green Supermix (Biorad). To test the correlation between circular RNAs candidates and the number of CAG repeats to faithfully recapitulate the bioinformatic analysis, we considered the expression of both linear and circular isoforms of the selected transcripts. We designed convergent and divergent primer sets for each candidate (Supplemental_Table_S6). To specifically target the circular isoform, the backspliced junction was selected. Expression values were obtained employing the 2^^-ΔΔ Ct^ method, normalizing with *Polr2a* and *Actb* housekeeping genes (HKG) and relativizing to Q111. The selected HKGs were chosen among a panel of 5, showing a more stable expression in our *Htt* KI samples. PCR amplicons were loaded on a 2% agarose gel, bands were carefully excised, purified using PureLink Quick Gel Extraction and PCR Purification Combo Kit” (Bioline) and eventually Sanger sequenced to finally prove their identity and the back-splice junction. GraphPad Prism (GraphPad Softwares) was used to establish a regression line between circular to linear ratios and the CAGs serie (20, 50, 89, 111). Finally, R^2^ values and corresponding p-values were calculated.

### Library preparation and RNA sequencing

For total RNA, samples were subjected to DNAse I treatment (Ambion) according to manufacturer instructions. Ribodepletion and barcoded stranded RNA-seq libraries preparation was performed by the European Molecular Biology Laboratory Genomic Core facility, following the standard Illumina protocols. Libraries were then sequenced with NextSeq 500 paired-end, 75bp reads, obtaining ∼60M reads/library. Similarly, miRNA libraries were generated by the European Molecular Biology Laboratory Genomic Core facility according to standard procedures and were sequenced on the Illumina HiSeq 2500 using single-end, 50bp reads obtaining ∼15M reads/sample.

### RNA-seq data, miRNA-seq and circRNA data analysis

Data from RNA-seq and miRNA-seq on mESC and mNPC allelic series were processed as follows. Reads filtering (minimum base quality of Q30) and adapter trimming was performed with Trim Galore (http://www.bioinformatics.babraham.ac.uk/projects/trim_galore/). Remaining reads were then aligned with STAR v2.5.3a (Dobin et al. 2013) to the mm10 assembly of the mouse genome coupled to the M14 Gencode genes annotation (10.1093/nar/gky955). STAR parameters were set as recommended by the DCC pipeline (doi: 10.1093/bioinformatics/btv656) to allow for the subsequent detection of circRNAs through chimeric junctions. Reads derived from circular RNA species were extracted from the WT, KO, and Q20 - Q111 samples of mESC and NPC cells by the DCC pipeline v0.4.4 (Cheng et al. 2016), run with default parameters. Repeats annotation provided to DCC was obtained from the UCSC Genome Browser (doi: 10.1093/nar/gkx1020), by combining the RepeatMasker and Simple Repeats track for the mm10 mouse assembly. Reads from circRNA host genes were also quantified by DCC in the same run. Total transcript mass was computed for a given transcript as the sum of its linear and corresponding circular RNA reads count. The relative abundance of the circular form of the transcripts was then computed as circular RNA read counts / circular + linear RNA read counts.

All p-values were computed by a Wilcoxon rank-signed test unless explicitly stated. Exons/introns annotation and the number of protein-coding genes per chromosome were obtained from UCSC (doi: 10.1093/nar/gkx1020) for the mm10 mouse genome assembly. These annotations were then used to compute circRNAs spliced length and the enrichment of expressed circRNAs in chromosomes (by a Fisher test).

Gene Ontology and pathways enrichment were computed with the topGO (Alexa and Rahnenfuhrer 2021) and clusterProfiler (Yu et al. 2012) R packages, using a BH adjusted p-value threshold of 0.05.

Differential expression of circRNAs was computed by the CircTest R package (Cheng et al. 2016). circRNAs were filtered by requiring at least five reads in at least one sample and at least 5% of total transcript mass in at least one condition. circRNAs having an FDR lower or equal to 0.05 were considered to be differentially expressed.

### Motif analysis

The sequence of the 100nts upstream and downstream of significant skipped exons events were extracted with the biomaRt R package (Durinck et al. 2009). A motif search was performed with DREME v4.10 (Bailey 2011), considering only same-strand matches, using a background sequence set generated by DREME via shuffling of input sequences and a 1.0E- 03 threshold on the E-value to select significant motifs.

### Differential Expression and Differential Linear Alternative Splicing Analysis

For the *in vivo* differential splicing analysis, the raw data from the study of Langfelder P et al (2016) was used. Striatum (GSE65774), cortex (GSE65770) and liver (GSE65772) mRNA expression profile datasets were retrieved for the analysis through the online database HDinHD portal (https://www.hdinhd.org/). At each of 3 time points (2, 6, 10 months), 8 heterozygous knock-in mice from each of the 6 Htt CAG repeat lengths (Q20, Q80, Q92, Q111, Q140, and Q175) were used, resulting in 48 samples from each tissue and each time point. Raw reads were subjected to sequence quality control using FastQC (http://www.bioinformatics.babraham.ac.uk/projects/fastqc/). Removal of low-quality reads and trimming of the adapter sequences were achieved by Trim Galore (http://www.bioinformatics.babraham.ac.uk/projects/trim_galore/). In order to eliminate variability between sequencing runs the reads were trimmed to 45 bp by Trim Galore, resulting in the removal of only 4% of the reads. Raw sequences were aligned to mm10 mouse genome assembly (UCSC) with STAR RNA-seq aligner version 2.5.3a (Dobin et al. 2013) using standard settings. The differential alternative splicing (AS) events between each of 5 samples for *Htt* CAG repeat lengths (Q80, Q92, Q111, Q140 and Q175) and Q20 (as control) were identified by rMATS v4.0.1 (Shen et al. 2014) (http://rnaseq-mats.sourceforge.net) that detects five major types of AS events from RNA-Seq data with replicates (Shen et al. 2012, 2014). Analyses results were then further processed with R/Bioconductor.

For the *in-vitro* differential splicing analysis, both mESC and mNPC samples were subjected to the same pipelines used for *in-vivo* differential splicing analysis with same parameters and tools. For the analysis of the differential alternative splicing (AS) events of 3 samples for *Htt* CAG repeat lengths (Q50, Q92, and Q111) Q20 samples were used as control. For the analysis of the differential alternative splicing (AS) events in knockout cells, wild type samples were used as control.

Differential expression analysis was performed on striatum samples for *in vivo* with read counts output of the alignment which was performed in previous step. edgeR (v3.24.3) was used within the Bioconductor environment in R and p-values were adjusted for multiple comparisons using the Benjamini–Hochberg method within each contrast and genes with FDR-adjusted p-value < 0.05 were considered significantly differentially expressed. To identify the genes both differentially expressed and alternatively spliced (all the different AS subtypes were included), differential expression and alternative splicing results were overlapped.

The functional enrichment analysis of the genes with AS events was performed using the DAVID functional enrichment tool v6.8 mainly based on GO terms (biological process, cellular compartment and molecular function), KEGG pathway, InterPro and UniProtKB keywords. The enriched terms were filtered according to FDR adjusted P-value <0.05.

## DATA ACCESS

The RNA-sequencing runs for mESC and NPC lines were deposited in GEO with ID GSE175658.

## ACKNOWLEDGEMENTS

We are grateful to the members of the Biagioli’s laboratory for helpful discussions and support during time. We wish to thank Vladimir Benes and the EMBL Genomics Core Facilities, Heidelberg, Germany for assistance during libraries preparation and sequencing. This work was supported by the University of Trento, the Huntington Society of Canada’s NEW PATHWAYS award, the HDSA Human Biology Project and the EHDN 1041 to MB and the National Institutes of Health (NS049206) to V.C.W. MB was a recipient of a Marie Skłodowska Curie reintegration fellowship (The European Union’s Horizon 2020 Research and Innovation Program) under the grant agreement No. 706567.

## AUTHORS’ CONTRIBUTION

D.A. was responsible for differential expression and linear alternative splicing (AS) analyses. A.M. and T.T. conducted most of the wet lab work, including mESCs differentiation to mNPCs, RNA collection and extraction, reverse transcription and PCR. J.D. and E.D. provided experimental validation to linear AS differences. L.D. and A.M. validated circRNAs expression differences. G.B. and M.K. contributed to mouse tissues collection and animal handling and caring. F.D.L. helped with RNA collection and initial RNA libraries preparation. J.Z. and L.C. supervised the differentiation of mESCs to mNPCs. V.C.W. provided mouse striatal tissues for AS validation. C.D. provided computational methods and suggestions for circRNAs data analysis. G.B. contributed to figures’ design. E.K. assisted with initial quality controls of the sequencing data. S.P. supervised D.A. during RNA-seq and AS analyses. E.D. conducted miRNA-seq and circRNAs data analysis. D.A., E.D., S.P., provided critical advice on statistical analyses. M.B., E.D., S.P. conceptualized and designed the study. D.A., J.D., E.D., G.B. and M.B. wrote the manuscript. M.B., E.D., S.P. supervised the project. All authors revised, read and approved the submitted version.

